# Identifying cancer cells from calling single-nucleotide variants in scRNA-seq data

**DOI:** 10.1101/2024.02.21.581377

**Authors:** Valérie Marot-Lassauzaie, Sergi Beneyto-Calabuig, Benedikt Obermayer, Lars Velten, Dieter Beule, Laleh Haghverdi

## Abstract

Single cell RNA sequencing (scRNA-seq) data is widely used to study cancer cell states and their heterogeneity. However, the tumour microenvironment is usually a mixture of healthy and cancerous cells and it can be difficult to fully separate these two populations based on transcriptomics alone. If available, somatic single nucleotide variants (SNVs) observed in the scRNA-seq data could be used to identify the cancer population. However, calling somatic SNVs in scRNA-seq data is a challenging task, as most variants seen in the short read data are not somatic, but can instead be germline variants, RNA edits or transcription, sequencing or processing errors. Additionally, only variants present in actively transcribed regions for each individual cell will be seen in the data. To address these challenges, we develop CCLONE (Cancer Cell Labelling On Noisy Expression), an interpretable tool adapted to handle the uncertainty and sparsity of SNVs called from scRNA-seq data. CCLONE jointly identifies cancer clonal populations, and their associated variants. We apply CCLONE on two acute myeloid leukaemia datasets and one lung adenocarcinoma dataset and show that CCLONE captures both genetic clones and somatic events for multiple patients. These results show how CCLONE can be used to gather insight into the course of the disease and the origin of cancer cells in scRNA-seq data.

## 1 Introduction

Cancer is a multistep process driven by somatic mutations in which healthy cells progressively evolve into cancerous states. Quantifying how cancer cell states differ from healthy states helps us understand this disease and provides potential therapeutic targets. Single cell RNA sequencing (scRNA-seq) has emerged as a powerful tool to study cancer cell states and their heterogeneity. However, the sampled tumour microenvironment is usually a mixture of healthy and cancerous cells [1], and fully separating these two populations can be difficult based on transcriptomics alone. Measuring mutational status and gene expression in the same single cell would allow us to more accurately identify the cancer population through mutations and relate that information to the observed transcriptional states. However, sequencing the entire genome and transcriptome of the same individual cell is costly and has low throughput [2–4]. Another option is to sequence only targeted genetic regions containing somatic mutations alongside the transcriptome [5–8], but this requires prior knowledge on the position of these mutations in each sample. These positions are usually inferred from preceding bulk DNA sequencing or through panel testing of known cancer associated genes. These current approaches require an adapted experimental design and carry additional costs and delays, which highlights the need for computational methods that can automatically and at large-scale, be applied to the prevalent uni-modal scRNA-seq datasets.

In the past, large copy number variants (CNVs) have been used to identify cancer populations in scRNA-seq data [9–11]. Since in that case the mutations cover large areas of the genome, the tools can leverage the read data across multiple adjacent regions to identify the CNVs. Mitochondrial variants (MVs) with high heteroplasmy have also proven helpful to study the cell lineage in cancer and healthy tissue due to the high mutation rate of the mitochondrial genome, large number of mitochondria per cell, and strong expression of mitochondrial genes [12–14]. However, not all cancers have CNVs or high heteroplasmy MVs and these events are found at different frequencies in different cancer types. In the absence of CNVs and usable MVs, recovery of cell lineages from scRNA-seq data has not been addressed up to date.

Single nucleotide variants (SNVs) observed directly in scRNA-seq reads could help identify cancer cells. However, calling SNVs confidently from scRNA-seq data is a challenging task. First, only variants present in actively transcribed regions will be seen in the data. Even then, we might not catch these variants in every cell due to the low average coverage of individual positions and of allelic dropout. Furthermore, when calling variants against the human reference genome, most variants seen in scRNA-seq reads are not somatic mutations but can be germline variants, RNA edits or transcription, sequencing or processing errors. In other words we will completely or partially miss most somatic SNVs, and most identified SNVs will not be somatic. Because of the high uncertainty of this data, SNV calls from scRNA-seq data are often very strictly filtered to ensure that mostly true somatic variants remain to identify the cancer cells [15]. This strict filtering can come at the cost of the identification of cancer cells in several lineages if they do not contain well covered high-confidence somatic variants. We hypothesised that by using methods which account for the uncertainty in the data, we could incorporate more variants (including low confidence variants) for identification of the clonal structure.

In this work, we show that SNVs called directly from scRNA-seq data can be used to identify cancer clonal populations. We introduce CCLONE (Cancer Cell Labelling On Noisy Expression), a tool adapted to handle the uncertainty in this big data. CCLONE is a fully automated tool that can be applied on new or existing scRNA-seq samples. We validate our results on three single-cell datasets with known cell clonal identities inferred based on targeted amplification of known SNVs, MVs and CNVs. The first two datasets present 19 patients with acute myeloid leukaemia (AML) [14,16], which is a cancer typically characterised by a low mutation load [17]. The third dataset presents 7 lung adenocarcinoma patients [18] which is typically characterised by a much higher mutational load [19]. For multiple patients, our method is able to reproduce the known clonal structure without using any prior information on the samples. The method also returns a set of SNVs enriched in each identified clone along with their expected allele frequency (i.e., homo/hetero-zygocity) in each clone. We show that these variants enhance interpretability of the results and points to real somatic events which inform about the course of disease evolution and the origin of cancer cells.

## 2 Results

In this section we first introduce and describe the CCLONE workflow. We then validate the tool on the AML and lung adenocarcinoma datasets, and show examples where the method helps get a better understanding of the analysed samples. Lastly, we show how the method’s likelihood of success depends on the data quality and the resulting capture of sufficient somatic variants.

### Overview of CCLONE workflow

For a fixed set of run parameters, the CCLONE workflow includes three main steps; i) variant calling and filtering, ii) clonal assignment using a weighted non-negative matrix factorisation, iii) statistical evaluation of the results. CCLONE runs this workflow for multiple sets of parameters and returns the most statistically favourable result as the final output.

As input, CCLONE takes annotated variant call data from scRNA-seq data (Figure 1.A). Cancer driver variants, passenger variants, or somatic variants found only in healthy clones can all be used to identify the clones. Ideally, the set of selected variants would contain as many somatic events as possible and as few non-somatic variants as possible. Therefore, assuming variant calling with no prior information on somatic events (such as we get from the bcftools variant calling pipeline [20]), CCLONE filters the likely non-somatic variants to maximise the signal-to-noise ratio (Figure 1.B). RNA edits, whose occurrence patterns are highly cell type specific [21], and thus a potential confounder in the data, are filtered out based on reference annotation in REDIdb [22]. Variants with very low coverage, or very low minor allele frequency (MAF) cannot reliably be leveraged to identify genetic clones and are also filtered out. Heterozygous germline variants are expected to be found equally in all cells. However, their occurrence can be associated to somatic events if they are located within regions either lost through a deletion or loss of heterozygosity event (LOH), or not expressed through strong imbalance in allelic expression (potentially due to a somatic variant in the regulatory region). Because of high variability between cancers, types of mutations and clone population size, a different filtering threshold might maximise the signal-to-noise ratio for different samples. To allow the model to make use of as many mutational events as possible, while excluding as many non-somatic variants as possible, we try different filtering thresholds and later allow the model to select the most informative set (Figure 1.B).

**Figure 1:**
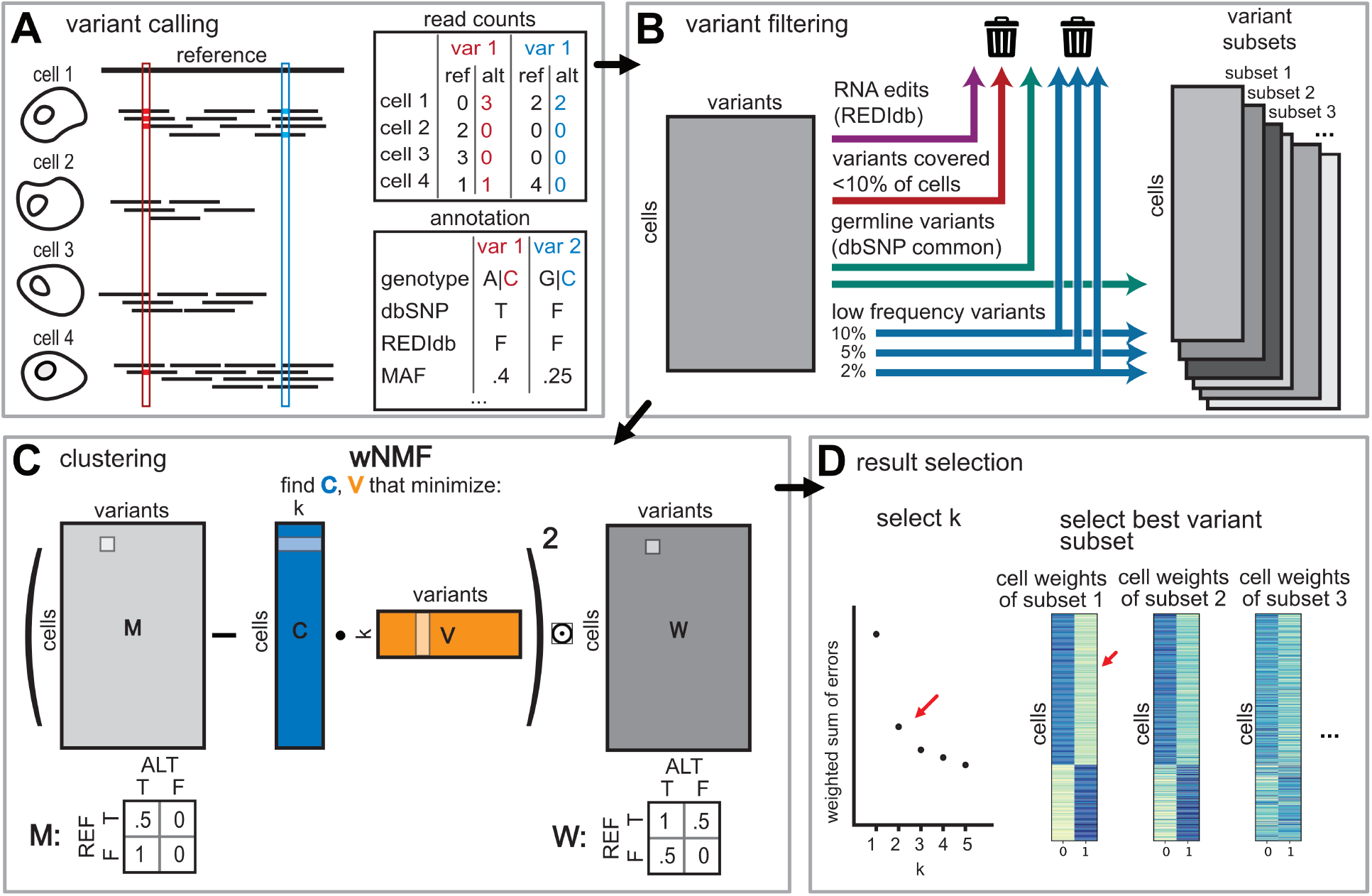
Overview of CCLONE workflow. (A) Variants are called from single cell short read data. The short reads differ from the reference in two positions (outlined in red and blue). For every variant, we keep the number of alternative and reference counts, and extract variant annotation. (B) We filter the variants based on database annotation as well as coverage and frequency. Because the best filtering threshold might differ between samples, we try different thresholds resulting in multiple variant subsets. (C) We use a weighted NMF to discover the hidden clonal structure in the variant call data. For this, the read count data is transformed into the observation matrix M corresponding to the discretized VAF, and a weight matrix W that reflects the confidence that we have in each value in M. The wNMF is calculated for every variant subset and a range of number of factors *K*. (D) To select the best result, we first select the best number of factors *K* according to the elbow method on the weighted sum of errors. To select the best variant subset, we compare the computed cell factors (matrix C) for every subset, and select the wNMF output with the largest (i.e., closest to 0 for negative values) orthogonality score *s* between the factors (Equation 4) as this reflects a clearer separation between the clones.

After filtering, we want to recover the hidden clonal structure from the cell-variant call matrix *M* (Methods 4.2.1.). We expect somatic variants to co-occur within genetic clones, but the variant call data is still both noisy and sparse. Non-negative matrix factorisation (NMF) has been widely used to capture hidden structure in noisy scRNA-seq data [23, 24]. However, NMF assumes that the data is complete, while we will only observe a variant if the position is covered, i.e. actively expressed in that cell. To account for this, we use instead a weighted NMF (wNMF), that uses a weight matrix *W* to reflect how confident we are in each value of the variants call matrix *M* (Figure 1.C, Methods 4.2.1.) based on each variants’ coverage in each cell. In wNMF, we try to learn the hidden factor matrices *C* (cell factors) and *V* (variant factors) that minimise the weighted sum of squared errors *E* of recovering the input data matrix *M* :

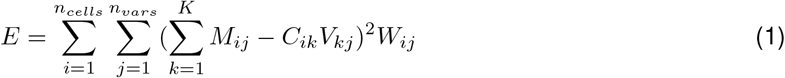

We allow the model to choose the number of factors *K* to reflect the number of genetic clones clearly distinguishable in our variant call data based on the elbow method on *E* (Figure 1.D, S1 shows the elbow on real data and Supplementary note A for discussion of the elbow method) [25].

We run the wNMF on all variant subsets and then allow the model to select the most informative set. Because the different subsets contain variants with different properties (different minimal MAF, inclusion / exclusion of germline variants), and thus different levels of expected uncertainty, the sum of squared error *E* is not directly comparable between the subsets. Here, we use instead the assumption that if the clones are clearly distinguishable in the variant data and captured by the model, then the cell factors reflecting these clones should be clearly separated i.e. uncorrelated. We select the best result based on the orthogonality score of the cell factors (Figure 1.D, Supplementary note B).

If the model succeeds, the cell factors should give us the final clonal assignments of that patient. The variant factors reflect the variants enriched in these clones and can be used to find disease relevant mutational events. The factors are not directly assigned a label of healthy or cancer, but must then be labelled through prior knowledge. In this work, we use the presence or absence of either known healthy or cancer cell types in each cell factor to label the factors (Supplementary note C). We consider the model to be successful if the factors can be clearly labelled as healthy and cancer. This ensures that the captured factors correspond to cancer and healthy clones, and that the separation between the two is well defined in the data. Alternatives to this approach are discussed in Supplementary note D.

### Application on AML patients datasets

We validate CCLONE on two AML single cell datasets. AML is usually characterised by a low mutation load. These mutations cause a block in differentiation of the hematopoietic stem cells (HSCs) resulting in the malignant expansion of aberrant progenitor cells called “blasts “[26]. This population is fuelled by leukemic stem cells (LSCs) that are transcriptionally similar to normal HSCs and difficult to target. Because of the low mutation load and presence of difficult to identify LSCs, AML provides a good test case on which to validate our method. The first analysed dataset contains 4 patients sequenced with SmartSeq2 [14], and the second dataset contains 15 patients sequenced with 10X [16] (Figure S2 shows the cell types and patient labels on UMAP). In both of these datasets, the cell labels as healthy or cancer could previously be recovered in some patients (respectively 2 and 11 patients) based on MVs and targeted amplification of known somatic SNVs in [14] and additionally through CNV in [16]. The two additional Smart-Seq2 patients had partial cell labels based on a single nuclear somatic variant. We call these previously recovered cancer and healthy cell labels “reference labels “in the following sections. The two datasets also allow us to compare the success rate with data from different sequencing technologies.

### CCLONE identifies cancer cells

CCLONE successfully identified cancer cells in all four SmartSeq2 patients (patients P1-P4 in Figure 2.A) and seven of the 10X patients (patients A1, A2, A6, A7 and A13-A15 in Figure 2.B). We use the T cells and Blasts to label the factors (Supplementary note C). Cells with similar weights for the healthy and cancer factors (difference in weights *<* .3) are labelled as undetermined. The method clearly groups the aberrant progenitors and the T cells in separate factors as expected for genetic clones in AML, while the stem cell populations are a mixture of healthy and cancer cells. We compare the labels for the patients where we have both reference cell labels and new labels (Figure 2.C.). We find a nearly complete overlap between the labels for patients A1, A2 and P2 (Figure 2.C.).

**Figure 2:**
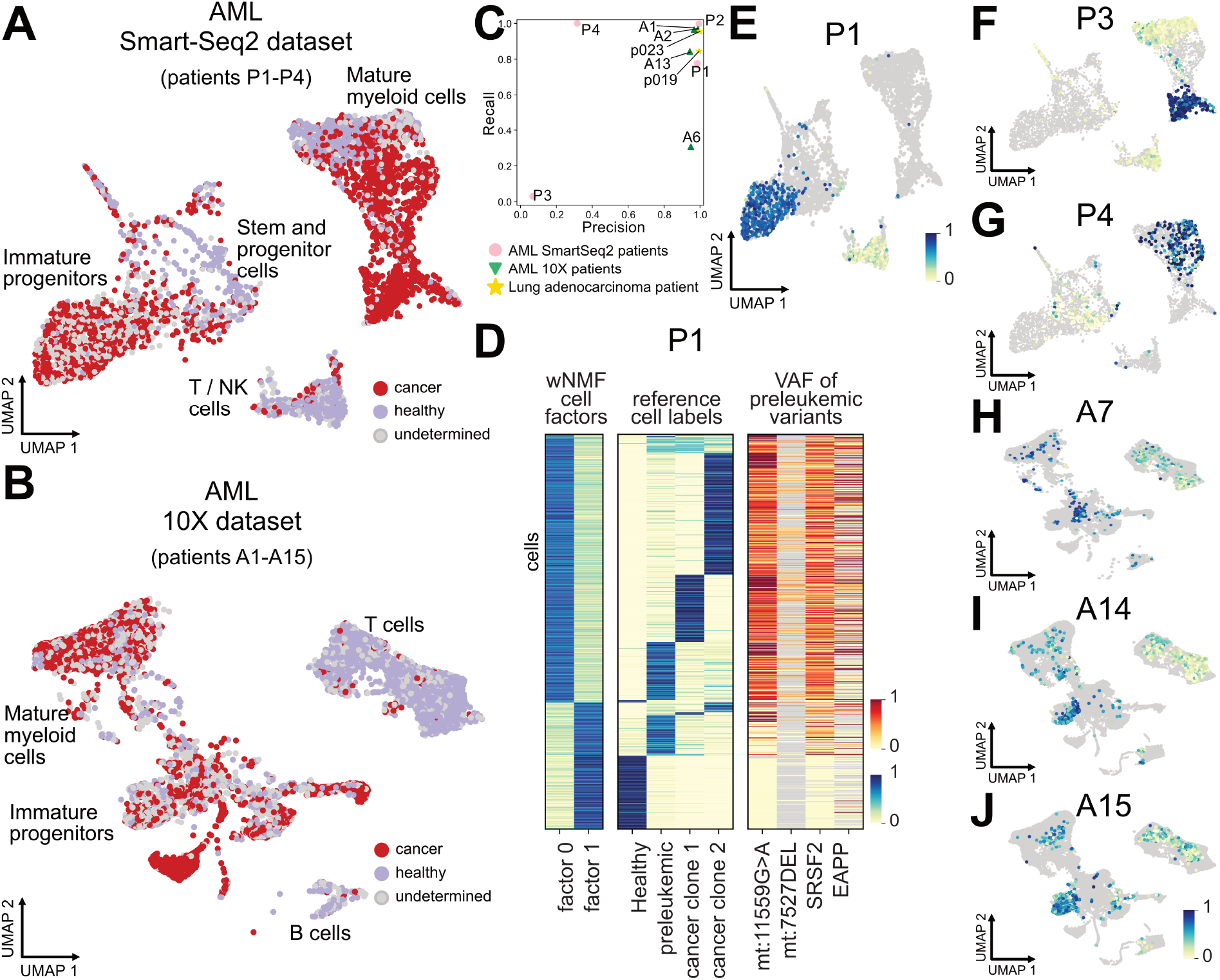
Cell factors capture genetic clones. (A-B) Cell assignments to healthy or cancer based on the wNMF cell factors plotted on the UMAP for the AML Smart-Seq2 dataset [14] in A and the AML 10X dataset [16] in B. (C) When present, we compare our cancer cell labels to the reference cancer cell labels from each dataset, and return the precision and recall for every patient. Note that reference cell labels are not available for all patients. (D) Heatmap comparing the wNMF cell factors to the reference cell labels and to the VAF of the variants used to separate the preleukemic population from the healthy one. (E) Cancer cell factor 1 of patient P1 coloured on the UMAP. Cells from patients P2-P4 are shown in grey for ease of comparison. (F-J) Cancer cell factors for patient P3, P4, A7, A14 and A15 coloured on the UMAP. Cells from the other patients of the dataset are shown in grey for ease of comparison.

For patient P1, A13 and A6, we are identifying a subset of the cells labelled as cancer in the reference. As cancer is continuously evolving through acquisition of new mutations, we likely have multiple cancer clones present in parallel in the sample. It is possible that our method identifies a subclone of the reference, as shown in the example of patient P1 (Figure 2.D and E). Here, the reference differentiates healthy cells, and the cancer population containing the preleukemic clone and two cancer subclones. Comparing the wNMF cell factors to the reference labels, we see that the reference healthy cells are predominantly assigned to factor 1, while the two cancer clones are assigned to factor 0. The preleukemic population I split between the two factors. The preleukemic cells assigned to factor 1 have much lower variant allele frequency (VAF) of the two MVs mt:7527DEL and mt:1159G>A than the other preleukemic and leukemic cells. This indicated that the preleukemic population might in reality correspond to two distinct subclones.

For patient P3 and P4 the reference is based on a single low-coverage nuclear variant each. For P3 there is no agreement between the reference and new labels. The reference labels are based only on a single IDH2 variant that is present in a subset of the mature myeloid cells. This variant is depleted in the subset of these cells that we identify as cancer cells (Figure 2.F and S3). This points towards there being two distinct genetic clones, each of which is captured by one approach. The IDH2 mutated cells are transcriptionally similar to monocytes, which could be either healthy or cancer. However, the population that we identify as cancer is transcriptionally very aberrant (for example expressing HBZ), indicating that they must be cancer cells that were missed previously. For patient P4 (Figure 2.G), the reference is based on a single very low coverage variant, and likely missed in many cancer cells (Figure S4). Here, CCLONE provides more complete cancer cell labels and potentially now captures all cancer cells instead of only a subset.

For patients A7, A14 and A15 we have no reference cell labels, but CCLONE could still identify a cancer population based on SNVs (Figures 2.H-J). These patients had no well-covered known leukemic SNVs and additionally no usable CNVs or MVs. This highlights the advantages of using a method that does not rely on prior knowledge of existing somatic SNVs, and consequently is not restricted to a small subset of the observed SNVs. Patients P3 and P4 are other examples where previous methods based on MVs and known leukemic SNVs are not sufficient to fully label the cancer cells. We further validate the clones based on known cancer and healthy cells within clones (Figure S5).

In this work, we do not use the MVs, nor directly call the CNVs for use with CCLONE. This means that we are potentially using a different set of somatic events than the reference to characterise the cancer cells. This can explain why we do not find the same exact same clones as the reference if the different variant subsets are found in different subclones.

### CCLONE finds cancer associated variants

On top of the cell factors, the wNMF also returns variant factors. To find clone-associated variants, we extract the variants with the largest difference in weight between the factors (>.3) and covered in at least 20% of the cells assigned to each clone. For the AML Smart-Seq2 patients, we have a whole exome (WE) cancer and control sample which we can use to validate and understand the variant factors.

The clone-associated variants of patient P1 are shown in Figure 3.A. Four of the SNVs associated with the cancer factor are supported by the WE, indicating that CCLONE is capturing somatic events. For P1 these four variants are also the only variants enriched in the WE cancer (Figure S6) and with sufficient coverage in the single cell data. A lot of the variants with high difference in weight between the clones are not found in WE, indicating that these could either correspond to variants characterising small subclones or some other genomic signal manifested in the noisy and imperfectly filtered variant calls. The VAFs of these variants show a very clear difference between the clones, and they co-occur with the known somatic SNVs, thus they can also be used by the wNMF to identify the clones. Patient P4 shows similar patterns to patient P1 (Figure S4).

**Figure 3:**
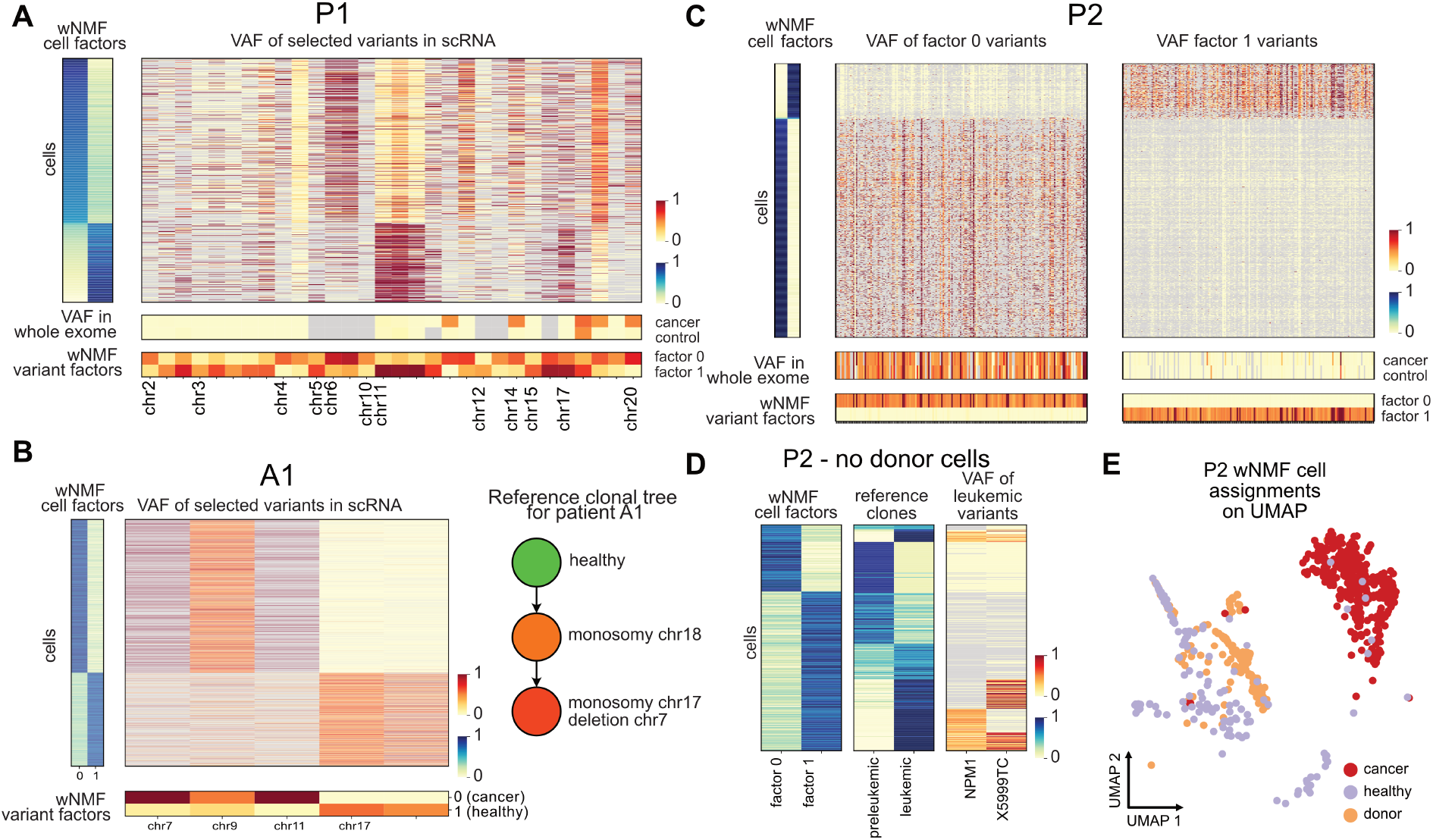
Variant factors capture somatic mutation events. (A) VAF for selected variants for the cells of patient P1. The cells are sorted by cell factors and the subset of variants are selected based on difference of weight between the variant factors (*>*.3). Grey values have too low coverage (*≤*2 reads for scRNA and *≤*5 for whole exome data). (B) VAF of selected variants (difference of weight *>* .3) for patient A1. The reference clonal tree was extracted from [16], and is based on their method CloneTracer. (C) VAF of selected variants (difference of weight *>* .3) for patient P2. The enriched variants in each factor are shown on two separate heatmaps for ease of visualisation of the different frequencies of observing these variants in the WE samples. Factor 0 (variants shown on right heatmap) corresponds to donor cells, and the variants characterising these cells are almost never found in the whole exome cancer or control, compared to the variants found in factor 1. (D) After excluding the donor cells and recalculating the wNMF, we compare the wNMF cell factors to the reference cell labels and to the VAF of the variants used to identify the leukemic population for P2. (E) Final CCLONE cell assignments for P2.

For some patients, we observe that variants observed at VAF close to 0.5 in the healthy populations are either lost (VAF *≈* 0) or fixated (VAF *≈* 1) in the cancer population (Patient A1, in Figure 3.B, A2 in Figure S7 and P3 in Figure S3). In these cases the method could be capturing CNV deletions or LOH, resulting in the loss of one allele and the heterozygosity of the germline SNVs overlapping that region. Figure 3.B shows the example of patient A1, where the monosomy on chromosome 17 was also captured by the wNMF. The other events on chromosome 7 and 11 point to further losses in those regions, one of which could correspond to the known deletion on chromosome 7. Another example of a patient where the wNMF potentially captures CNVs is patient P3 (Figure S3). The variants separating the cancer subclone from the other cells are predominantly germline variants (found in both WE cancer and control), and variants heterozygous in the healthy population are again either lost of fixated in the subclone.

For patient P2 (Figure 3.C), we find a very high number of variants associated with each clone, and those associated with clone 1 are found in both WE samples while those associated with clone 0 are found in none. Both of these variants subsets are enriched in known germline SNVs (dbSNP common). The extremely high number of likely germline variants clearly separating both factors point towards this patient potentially having cellular mosaicism. Such blood microchimerism can arise naturally if the patient had a twin through exchange of hematopoietic stem cells in utero, or after pregnancy [27, 28]. This was overlooked in previous analysis, and the reference cell labels reflect the separation between “donor “(factor 0) and patient cells (factor 1). Here, considering both cell factors and variant factors jointly allows us to get a more complete picture of the data. Excluding the donor cells, and rerunning variant filtering and wNMF, we find two clones, corresponding in parts to the separation between preleukemic and leukemic in the original publication [14] (Figure 3.D). The final labels for patient P2 are shown on the UMAP in Figure 3.E. The main disagreement between the reference preleukemic and leukemic labels and our assignments are in cells with low or no coverage for the leukemic variants used to label the reference (Figure 3.D), indicating that CCLONE is helping us refine the labels and now identifies all leukemic cells. This separation between healthy and cancer for P2 could only be identified after excluding the donor cells and the associate variant, as the signal-to-noise ratio for this pattern was otherwise too low. This highlights the importance of variant set selection.

The VAF plots of selected variants for all additional AML 10X patients are shown in Figure S7, and for patients P2 excluding donor cells in Figure S8.

### Data quality and mutation load determine success

CCLONE does not succeed in recovering clonal structure for all AML patients. The model needs multiple co-occurring somatic events to identify the clones, and these might not be found at sufficient frequency for all the patients. In particular, we see a much higher rate of success for the Smart-Seq2 patients, than for the 10X ones. This is likely due to the higher sequencing depth, and to the fact that Smart-Seq2 is not 3’ biased, resulting in a higher rate of capture for somatic events not located in the 3’ end. We thus capture a much higher number of variants for the Smart-Seq2 patients than for the 10X patients (Figure 4.A). Overall, we expect the method to work better at higher sequencing depths as we have higher odds of capturing somatic events across cells. The exception is given by patients A1 and A2 that are characterised by multiple large CNVs captured by the method. To simulate the success rate at lower sequencing depth, we subsample counts from our variant calls and rerun CCLONE. We then compare the results on the subsampled counts to the full counts in Figure 4.B. (S9 for recall and number of variants kept). As expected the precision significantly declines at lower simulated sequencing depths, although with high variability between patients, reflecting the variation in clarity of the signal.

**Figure 4:**
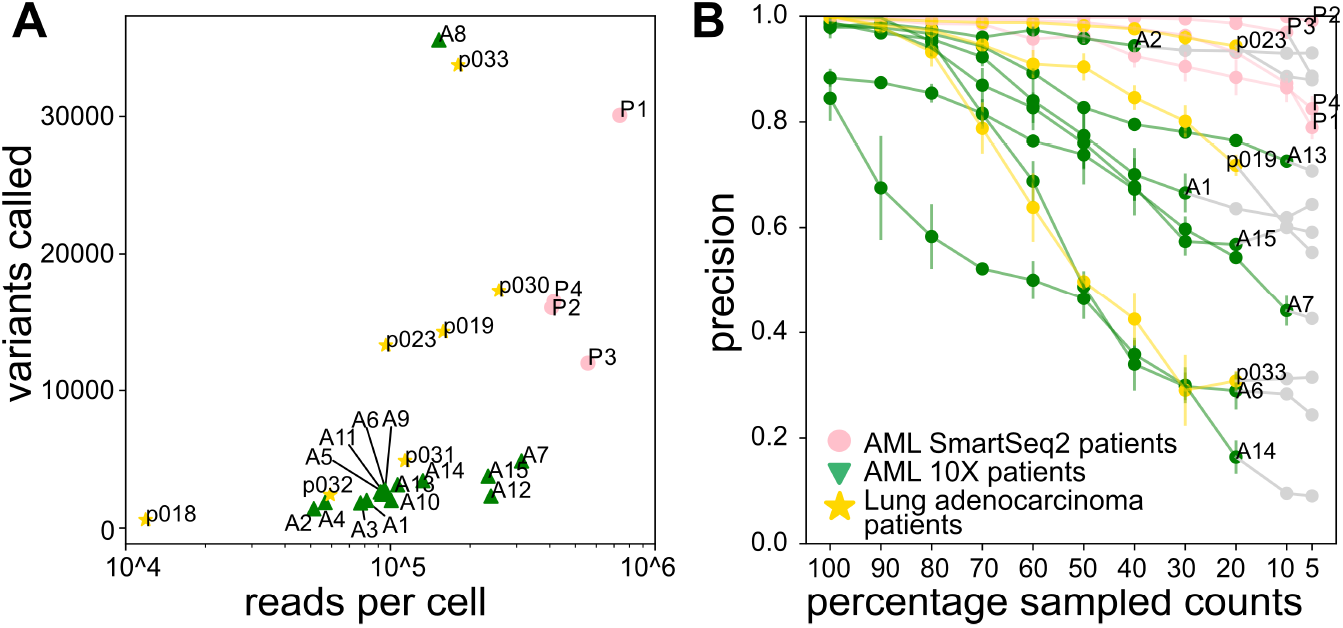
Method success is dependent on the quality of the data. (A) Total number of called variants as a function of the number of reads per cell for each patient. Here we include all called variants covered in 10% or more of the cells. Note the log scale on the x axis. (B) For every patient we randomly subsample counts from the reference and alternative count matrices, and rerun the wNMF on the subsets. Comparing the cancer cell labels between the full dataset and the subset, we get the precision as a function of the percentage of sampled counts. The values are plotted in grey if the number of variants with sufficient coverage is lower than 100.

In total, we recover cancer clones in 11 of 19 AML patients, even if the captured clone does not always cover all cancer cells (Figure S5). This is a success rate approaching the combined use of targeted amplification of known SNVs, MVs and CNVs (success for 15 patients). This success rate is particularly notable if we take into account the significantly higher ease of application of CCLONE to new and existing datasets.

As the success of the wNMF is correlated with the number of captured somatic events, we hypothesised that for tumours with higher mutation load we might still succeed in capturing the clones at low sequencing depth. To test this, we applied CCLONE on a 10X lung adenocarcinoma [18] with 1.5×10^4^-1.5×10^5^ reads per cell, shown in yellow in Figure 4.A. Here, the reference cancer cell labels are equivalent to the ones used in the original study and based on CNVs. The wNMF needs the presence of at least 2 genetic populations at sufficient frequencies to find patterns of variant co-occurrence. Because of this, we exclude patients that have almost only (>97%) tumour cell types, leaving us with 7 patients (Supplementary Figure S10). Figure S11 shows the cell types and patient labels on a UMAP. CCLONE succeeds in identifying the cancer population in 3 of the 7 analysed patients (Figure S12 shows the labels on the UMAP) and the cancer cell labels have a high overlap with the reference for p019 and p023, and correspond to a subset of the cells for p033 (in yellow on Figure 1C and S5). This success rate is lower than expected. Nonetheless, the method still succeeds for some patients, and the VAF plots of selected variants (Figure S13) helps us identify likely somatic events.

### Computational efficiency

The two most computational expensive steps of CCLONE are variant calling and wNMF. We report the runtime on a Dual Xeon E5-2650v2 (8cores/2.6GHz) and 15 GB of memory for variant calling with Cellsnp-lite in Figure S14, and for the wNMF in Figure S15. The runtime of both steps scales linearly with the input size.

For a 10X patient with 10000 cells, variant calling with Cellsnp-lite over all chromosomes in parallel, we estimate to take about 48 hours. Depending on the number of variants, subsequent analysis of the variant calls with CCLONE we estimate to take up to 12 hours.

### Data and software availability

The raw sequencing data of the two AML datasets are available at the European Genome-Phenome Archive with the accession ids EGAS00001003414 [14] and EGAS00001007078 [16]. All notebooks necessary to reproduce the results reported in this paper and the anonymised variant call data is available on our GitHub Page: https://github.com/ValerieMarot/clonal_tracing_notebooks. A python package of the CCLONE pipeline can be found at https://github.com/ValerieMarot/clonal_tracing_package.

## 3 Discussion

In this manuscript, we show that cancer cells can be identified from SNVs called directly from scRNA-seq data. These calls tend to contain many non-somatic variants and often have very low to no coverage in individual cells. We introduce CCLONE, a method adapted to work with uncertain variant calls. We validated the method on 2 AML datasets (19 patients) and a lung adenocarcinoma dataset, and show that the method captures genetic clones. The interpretable output of the method also allows us to find disease associated variants pointing to somatic events. By jointly considering all cells and all variants to find patterns of co-occurrence, CCLONE can identify patterns that might be missed by other methods. This is nicely exemplified by patient P2, where the cellular mosaicism was missed by the MutaSeq analysis, even though the same clones were identified. Another example is patient P3, where our approach finds a different cancer subclone that was overlooked in previous analysis.

Per default, CCLONE takes all variants called from scRNA-seq data as input, and automatically tries to find the most informative variant subset. This approach avoids more complicated and costly variant filtering procedures and makes the method easy to apply on new samples. Nevertheless, the set of variants used as input to the wNMF will influence its output. Including too many non-somatic variants causes a reduction of the signal-to-noise ratio, and might result in losing the clear separation of clones. On the other hand a too strict filtering can result in the exclusion of somatic events and loss of signal. Another issue can arise from the presence of correlated non-somatic variants such as cell type specific RNA edits. Here the wNMF would cluster according to these variants and the resulting clones will not reflect genetic information. This highlights the importance of sensible filtering criteria of the variants subset used as input. In the absence of prior knowledge (such as identified through panel testing of known disease genes or WE data), we propose to try different filtering thresholds for the variants used as input to the wNMF and afterwards determine the best result based on the metric introduced in Equation 4. Another solution would be to include only variants that are very likely of somatic origin, either through prior knowledge or through filtering and statistical testing [29, 30]. However, allowing the model to make use of likely germline variants if they are informative can result in the capture of CNVs, as was the case for patient A1. Furthermore, excluding all uncertain variants could come at the cost of resolution for cells that do not have coverage for the small subset of selected variants, as shown on the unreliable reference for patient P3 and P4.

CCLONE relies on finding groups of variants that tend to co-occur within genetic clones (i.e. the healthy or cancer populations). Therefore, the mutational load of the analysed sample, as well as the capture rate of these variants are crucial determinants of CCLONE’s success for a specific sample. The capture rate depends both on the mean sequencing depth and also on the properties and biases of the used sequencing technologies. As shown in section “Data quality and mutation load determines success “3’ biased technologies such as 10X will miss more variants than technologies with coverage over the full length of the transcript. For cancer samples with low mutational load, such as is expected for most AML patients, we recommend higher sequencing depth (*≈*1*e*6) and coverage over the full length of the transcript. For cancer types with very high mutational load, the co-occurring variants might still be found at lower sequencing depths with 3’ biased technologies, although very low sequencing depth can still result in missing the existing signal.

The wNMF step of CCLONE tries to find groups of co-occurring variants across cells in an unsupervised manner. The smaller the group of co-occurring variants and the smaller the clonal cell populations, the smaller the decrease in error E. One consequence of this, is that very small clones can be missed by the wNMF, as the decrease in E approaches background noise. To ensure that both the healthy and cancer populations have sufficient size to be captured by the wNMF, we restrict our analysis to patients that have both sufficient healthy and cancer cell types. We recommend each clonal population to be present at frequencies of at least 10% when running CCLONE on new data. Alternatively, prior knowledge during variant selection can also help enhance the signal-to-noise ratio to allow for the identification of smaller clones. In this work, we further use these cell types to annotate and validate the factors, highlighting again the need for sufficient cells of these cell types in the sample. Other prior knowledge on the analysed samples (such as known somatic events present in the data) could be used instead in the absence of this information.

Cancer is continuously evolving through acquisition of new somatic mutations and clonal expansion. Therefore there is an inherent uncertainty in any attempt to group the cells into separate groups of healthy and cancer cells. Depending on the time of acquisition of each somatic variant, these variants might be present in all, or in different subsets of the cancer cells. This imbalance between variants found in different subclones might be particularly pronounced if the cancer acquires a mutator phenotype, resulting in a higher mutation rate in the corresponding clone [1]. In the presence of multiple subclones with different somatic variants, the subset of variants used as input to the wNMF will then determine which of these we identify. As a result, for early cancer clones, the wNMF is not guaranteed to identify all cancer cells. This is exemplified by patients P1, P3 and A6, where we identify a subset of all cancer cells as the cancer population (Figure 2C and S4). For patients P1 and P3, we could show that the identified subset of cells corresponds to a genetically distinct population of cells, indicating that we are capturing a subclone of the cancer population.

CCLONE’s success rate (11 of 19 AML patients) in using SNVs seen in scRNA-seq data for identification of clonal structure in cancer samples motivates further computational developments in this direction. These types of approaches that require no previous knowledge about existing mutations can provide clonal insight into existing datasets, potentially avoiding the cost of additional experiments (such as WE sequencing) for samples where the clonal structure can be recovered from variants observed in the scRNA-seq data alone. In the future, one could consider other probabilistic approaches of handling uncertainty in the variant call data. These could include efficient ways of incorporating prior knowledge (e.g. through known somatic events) into semi-supervised classification algorithms. In this work we focused on extracting the information present in the uncertain SNVs and showing that this layer contains usable clonal information. Future methods could also combine the different information layers provided from nuclear SNVs, CNVs and MVs. This could help us get a more complete picture of the mutational journey of healthy cells towards cancerous states.

## 4 Methods

### 4.1 Processing of raw data

#### 4.1.1 Preparing the raw data and variant calling

The raw data was aligned to reference genome hg38 [31] with Star version 2.7.8a [32] for the MutaSeq data and Cell Ranger version 7.1.0 for the two 10X datasets. Variants were called single cells with Cellsnplite version 1.2.3 [33]. We annotate the variants with VEP [34] with custom annotation of common dbSNP germline variants (MAF*≥* 0.01 in at least one major population) [35] and RNA edits found in REDIdb [22].

#### 4.1.2 Variant filtering

To minimise potential artefacts in variant calling, we filter all variants found in repeat regions, according to RepeatMasker [36]. We filter all variants annotated as RNA edits. We remove all low coverage variants found with cov *≥* 2 in less than 10% of the cells.

To allow flexibility in filtering of germline variants and low coverage variants, we create 6 different variant subsets corresponding to the combination of different thresholds. These thresholds are exclusion / inclusion of germline variants and exclusion of variants with MAF of 2%, 5% or 10%. We then run the wNMF on each subset and later choose the most informative subset.

### 4.2 Clustering of cells and variants

#### 4.2.1 Input

The NMF takes as input an observation matrix *M* and a weight matrix *W* (both of dimension *n*_*cells*_, *n*_*vars*_). *M* is chosen to represent our best estimation of the true VAF, while *W* reflects the confidence we have in each value of *M*.

When working with nuclear variants, *M* contains the discretized (into three values 0, 0.5, 1 (corresponding to homozygous reference, heterozygous, and homozygous variant observations) VAF for every variant in every cells. We count an allele as observed, if we see at least two UMIs (or 2 reads for non-UMI technology) matching that allele, and an allelic frequency *≥* 0.05. We have:

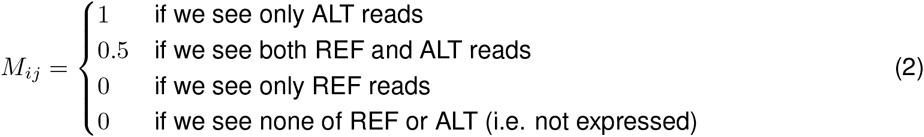

Observing only REF or ALT in a cell can either indicate that the cell does not have that variant, or it can be due to imbalances in allelic expression. To reflect this, we define *W* as:

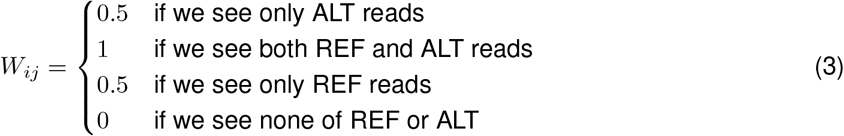

#### 4.2.2 wNMF

In NMF, we factorise the observation matrix *M*, into two matrices *C* of size (*n*_*obs*_, *K*) and *V* of size (*K, n*_*vars*_), with the constraint that these matrices have no negative elements. In wNMF, we further weight each value in *M* by its weight defined in *W* according to Equation 1.

Since the solution to Equation 1 is not unique, we find the optimal *C* and *V* matrices via an EM procedure; first *C* is (randomly) assigned and *V* is found by a non-negative least squares solver. Then *V* is fixed and *C* is solved analogously. We iterate these two steps until convergence (default of 1000 EM iterations). Figure S16 demonstrates the robustness of the final solution with respect to the initial (random) initialisation on real data.

Ideally, the number of latent factors *K* would reflect the number of co-occurring variant groups clearly identifiable from the data. If this information is known, the corresponding *K* can be used as input to the wNMF. In the absence of prior knowledge, we try to determine the best number of factors based on the elbow method on E (Equation 1). We run the wNMF for a range of *K* (default of 1 to 5), and get the error E for each of these. The decrease in E should be high while the new factors still capture bigger groups of co-occurring variants and level off when the new factors describe small groups or single variants. We use the kneedle algorithm [25] to automatically find the elbow. E as a function of *K* and the chosen elbow are shown in Figure S1 for all patients analysed in this work.

#### 4.2.3 Bootstrapping

The method to fit our wNMF does not guarantee finding a global minimum of the cost function, and the final factor matrices can vary over multiple rounds with random initialisation. To test its robustness with respect to noise in input data, we bootstrap the wNMF by randomly subsampling 90% of the variants and recomputing the factor matrices, with default of 10 bootstrap. We then align the results and get the mean and variance of the 10 bootstrap matrices. The mean *C* and *V* matrices gives us the final assignments, while the variance of the bootstrap matrices indicate how robust these assignments are over multiple bootstrap iterations.

#### 4.2.4 Selection of result

We run our wNMF on the different variant subsets, and select the best output as final result. Because the variant subsets include different number of variants which can have different properties (different types of event included, different MAF), the weighted sum of squared errors E is not directly comparable. Instead, we compare the cell factors *C* that have the same dimension for all subsets. We expect the cell factors to reflect our genetic clones, and for each cell to be predominantly assigned to one clone. Hence, the best result is chosen as the one with the largest (i.e., closest to 0 for negative values) orthogonality score *s* between the clones:

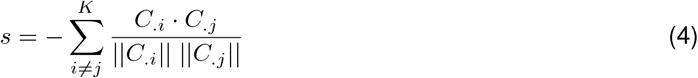

, where *C*_.*i*_ and *C.j* indicate the *i*-th and *j*-th columns of *C* respectively, and the nominator presents the inner product between them. The denominator presents the norm of the vectors.

## 5 Funding and Acknowledgements

Thanks to Philip Bischoff (Charité) for sharing the raw data of the lung adenocarcinoma cohort. Thanks to Melanie Fattohi (MDC) and Colin Cess (MDC) for proofreading. Thanks to Martin R. Siegert and to all IT of the MDC for maintenance of a high-quality HPC system and great support.

This study was supported by the Bundesministerium für Bildung und Forschung (BMBF) grant for ‘junior consortia in systems medicine’ to LH and LV (project number 01ZX1911). Additionally, LH and DB are supported by the Deutsche Forschungsgemeinschaft (DFG) SFB1588 grant (project number 493872418).

