## Supplementary notes and figures for "Identifying cancer cells from calling single-nucleotide variants in scRNA-seq data"

Supplementary notes and figures for
### "Identifying cancer cells from calling single-nucleotide variants in scRNA-seq data"

Valérie Marot-Lassauzaie[1,2,*], Sergi Beneyto-Calabuig[3,4], Benedikt Obermayer[5], Lars Velten[3,4], Dieter Beule[5,6], Laleh Haghverdi[1]

**1** Berlin Institute for Medical Systems Biology, Max Delbrück Center (BIMSB-MDC) in the Helmholtz Association, Berlin, Germany
**2** Charité – Universitätsmedizin Berlin, corporate member of Freie Universität Berlin and Humboldt-Universität zu Berlin, Berlin, Germany
**3** Centre for Genomic Regulation (CRG), The Barcelona Institute of Science and Technology, Dr. Aiguader 88, Barcelona 08003, Spain
**4** Universitat Pompeu Fabra (UPF), Barcelona, Spain
**5** Berlin Institute of Health at Charité – Universitätsmedizin Berlin, Core Unit Bioinformatics, 10117 Berlin, Germany
**6** Max-Delbrück-Center for Molecular Medicine in the Helmholtz Association (MDC), Berlin, Germany

*

**2 Supplementary figures** **5**

#### 1 Supplementary notes

##### Note A: Select number of factors K

The wNMF takes as input the observation matrix $M$, the weight matrix $W$ and the number of latent factors $K$.

To minimise the sum of squared error E, the wNMF finds groups of variants with correlated VAF patterns across cell groups (for example a set of variants that is seen only in the cancer population). These variants and cells can be summarised in a factor, resulting in a reduction of E proportional to the number of variants and cells expressing these variants in each group. The higher the number of co-ocurring variants, the higher the drop in E, which results in the first factors capturing the main axes of variant variation in the data. Once all groups of co-occurring variants are found, the remaining factors will capture smaller patterns, until the factors describe only individual variants.

Ideally, the inputed number of latent factors $K$ would reflect the number of clones present in the data. While in theory, the number of clones present in the sample should be well-defined, in practice, the number of clones that can be identified will depend on the captured variants. To ensure that the factors are not capturing background noise, we would like $K$ to reflect the number of co-occurring variant groups clearly identifiable from the data. If this information is known, the corresponding $K$ can be used as input to the wNMF. In the absence of prior knowledge, we try to determine the best number of factors based on the decrease in E. We run the wNMF for a range of $K$ (default of 1 to 5), and get the error E for each of these. The decrease in E should be high while the factors still capture bigger groups of co-occurring variants and level off when the factors describe small groups or single variants. E as a function of $k$ and the chosen elbow are shown in figure S1 for all patients analysed in this work. One consequence of this is that very small clones will be difficult to capture, as they cause only a small loss in E, maybe even smaller than the loss in E caused by strongly expressed individual variants.

#### Note B: Variant set selection for wNMF

As explained in the main text, it is important to maximise the signal-to-noise ratio in the variant set used as input for the wNMF. The cancer population size, and type of somatic events found (deletion of germline SNV, acquisition of somatic SNV) can greatly vary between samples. Because of this, the ideal variant filtering thresholds can also vary between patients. To account for this,SNV we try different thresholds and run the wNMF on each subset. Per default we try both to include and exclude known germline SNVs (based on common dbSNP variants [1]), and to vary the minimal MAF across cells between 2, 5, and 10. Because the properties of the included variants will vary between these subsets, the sum of squared errors E is not directly comparable. Instead, we use the assumption that if the model reliably captures the clonal structure in the data, we expect the cell factors reflecting these clones to be clearly separated. That is because we expect the cell factors to reflect our genetic clones, and for each cell to be predominantly assigned to one clone. A greater separation between the cell factors indicates that this subset has a clearer signal, and thus we assume that this is the best subset.

For a fixed number of latent factors $K$, and a set of variant subsets $V = \{V_0, \ldots, V_n\}$ with corresponding fitted wNMF cell factor matrices $\{C_0, \ldots, C_{V_n}\}$ all of equal shape ($n_{obs}$, $K$), we select the best subset $V_\star$ as the one that maximises the orthogonality score $s$ between the clones as defined in main text Equation 4.

We note that the subset of variants included as input to the wNMF will have a strong effect on the decrease in E. Both excluding and including too many variants can result in missing some / all clones, as either the signal is lost through exclusion of the relevant variants or the signal-to-noise ratio becomes too low by including too many non-relevant variants. This second case is exemplified by patient P2, where the separation between the healthy and cancer non-donor cells is missed until we exclude all variants characterising the donor cells.

#### Note C: Label latent factors

As output of the wNMF, we get the cell factor weights $C$ ($n_{obs}$, $K$), the variant factor weights $V$ ($K$, $n_{vars}$), as well as the orthogonality score $s$ as defined in main text Equation 4. The wNMF tries to find the biggest sets of co-occurring variants across cells, and summarises those by a factor. If the method succeeds we expect the factors to reflect genetic clones, i.e. one or multiple factors for healthy clone(s) and one or multiple factors for cancer clones(s). However, the wNMF does not directly label these factors as healthy or cancer.

In this work, we label the factors based on prior knowledge of cancer cell populations. For the AML samples, we know that the cancer does not give rise to either T and NK cells, and use those for labelling the clones. Clones depleted in these populations ($< 10\%$ of T / NK cells in that clone) are labelled as cancer clones, and clones containing these cell types are labelled as healthy. For patients with too few T/ NK cells (AML Smart-Seq2 patients P2 and P4), we use cancer cell types to label the clones. Clones depleted in Blasts ($< 10\%$) are labelled as healthy, and the others as cancer. For the lung dataset, the AT cells, ciliated cells and club cells are assumed to contain mostly healthy cells. Here too, clones depleted in these populations are labelled as cancer clones, and those that are not depleted in these cell types are labelled as healthy.

We also use these known healthy / cancer cell states to validate the clones. The wNMF tries to find groups of co-occurring variants across cells in an unsupervised manner. However, it is not guaranteed that such groups of variants are present in the data. Another problem could arise if the data contains co-occurring variants of non-somatic origin (for example missed RNA edits, or correlated artefacts). The sum of squared errors $E$ (main text Equation 1), reflects how well the present variation is captured by the wNMF. The orthogonality score $s$ (main text Equation 4), reflects how clear the separation between the cell factors is in the data, and thus the expected signal-to-noise ratio. Neither $s$, nor $E$ inform us whether the captured variation reflects genetic clones. To ensure that the variation captured by the wNMF corresponds to separation between healthy and cancer, we use instead the known cell types to validate the factors. If no factors are labelled as either healthy or cancer, we assume that the model has failed.

##### Note D: Alternatives approaches to label latent factors

In the absence of reference cell states (i.e. all cell states are mixture of healthy and cancer), prior knowledge on the variants used as input to the wNMF could be used for labelling and validation instead. If some variants correspond to known somatic events we could then verify that these have different weights between the factors, and use these for labelling of the factors. Another alternative would be to carefully curate the variant set used as input to the wNMF and ensure that only likely somatic events are used. By excluding all other potential sources of co-occurring variants, we would then ensure that the identified variation corresponds to somatic variation. However this exclusion of uncertain variants is potentially time consuming and error prone. It could also come at the cost of missing resolution for several lineages if they contain no well-covered curated variants.

In the absence of prior knowledge on the variants, a more careful analysis of the variants enriched in each factor could help in understanding, labelling and validating the factors. If we find multiple neighbouring variants found at VAF$\approx$ 0.5 in one population and either lost of fixated in the other population, they could point towards a deletion or LOH in that region (as shown for patient A1 in Figure 3, or P3 in Figure S3). Another option would be to look for potential driver variants in the enriched variants. This could be done by testing whether these variants are predicted to have an effect and are found within disease-associated genes. As shown in this work, the patterns of enriched variants can be very different between patients. Consequently, this approach would have to allow for flexibility and evaluating the enriched variants might be time consuming. It would also come at the cost of excluding patients with no identifiable somatic events in the enriched set.

#### 2 Supplementary figures

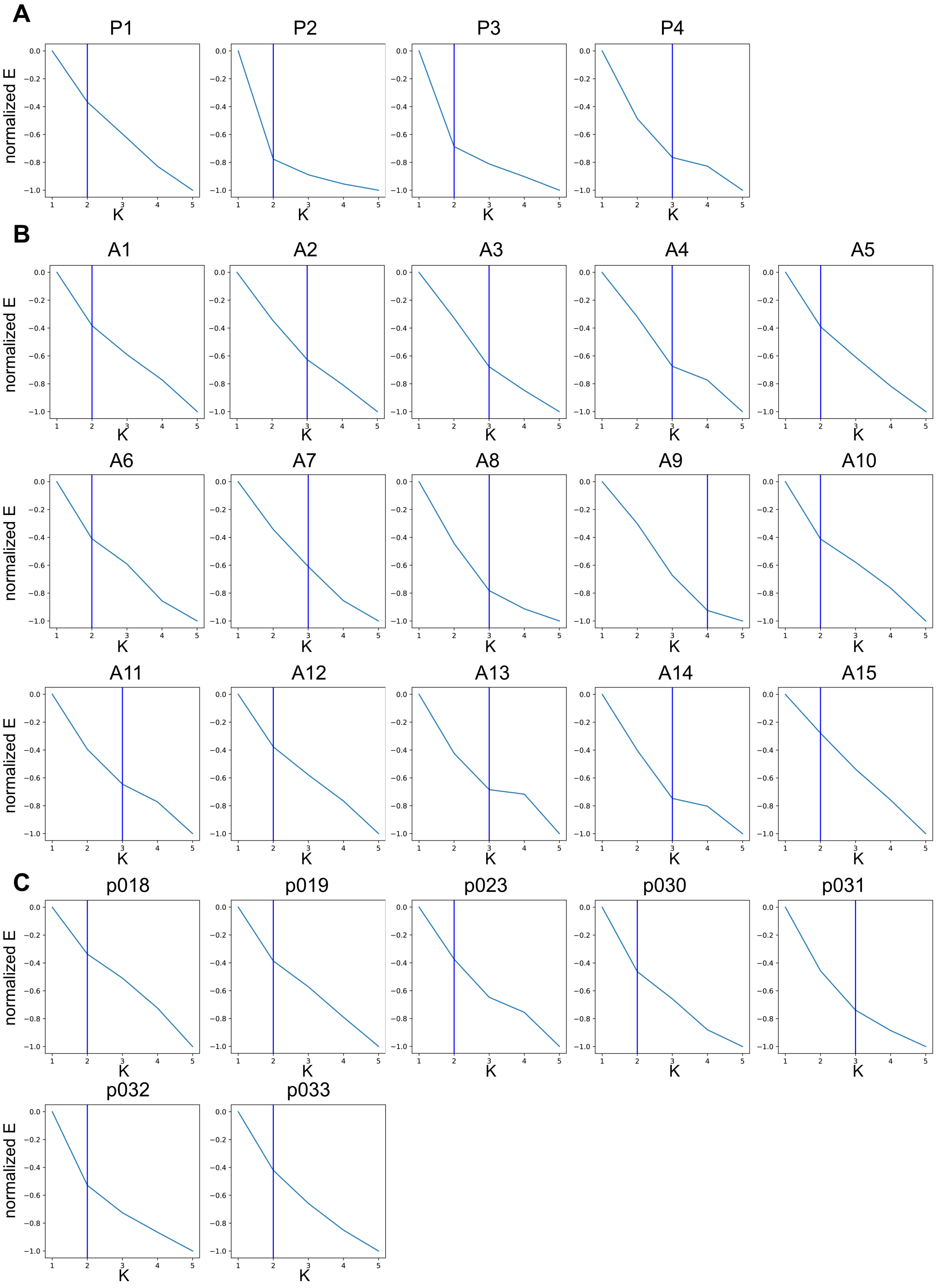

**Figure S1: Elbow plots for real datasets.** We show the elbow plots used to determine the number of clones on the the 4 AML Smart-Seq2 [2] patients in (A), on the 15 AML 10X patients in (B) [3] and on the 5 lung adenocarcinoma [4] patients in (C). We use the kneedle algorithm [5] to automatically find the elbow based on the normalized sum of errors E (Main text Equation 1).

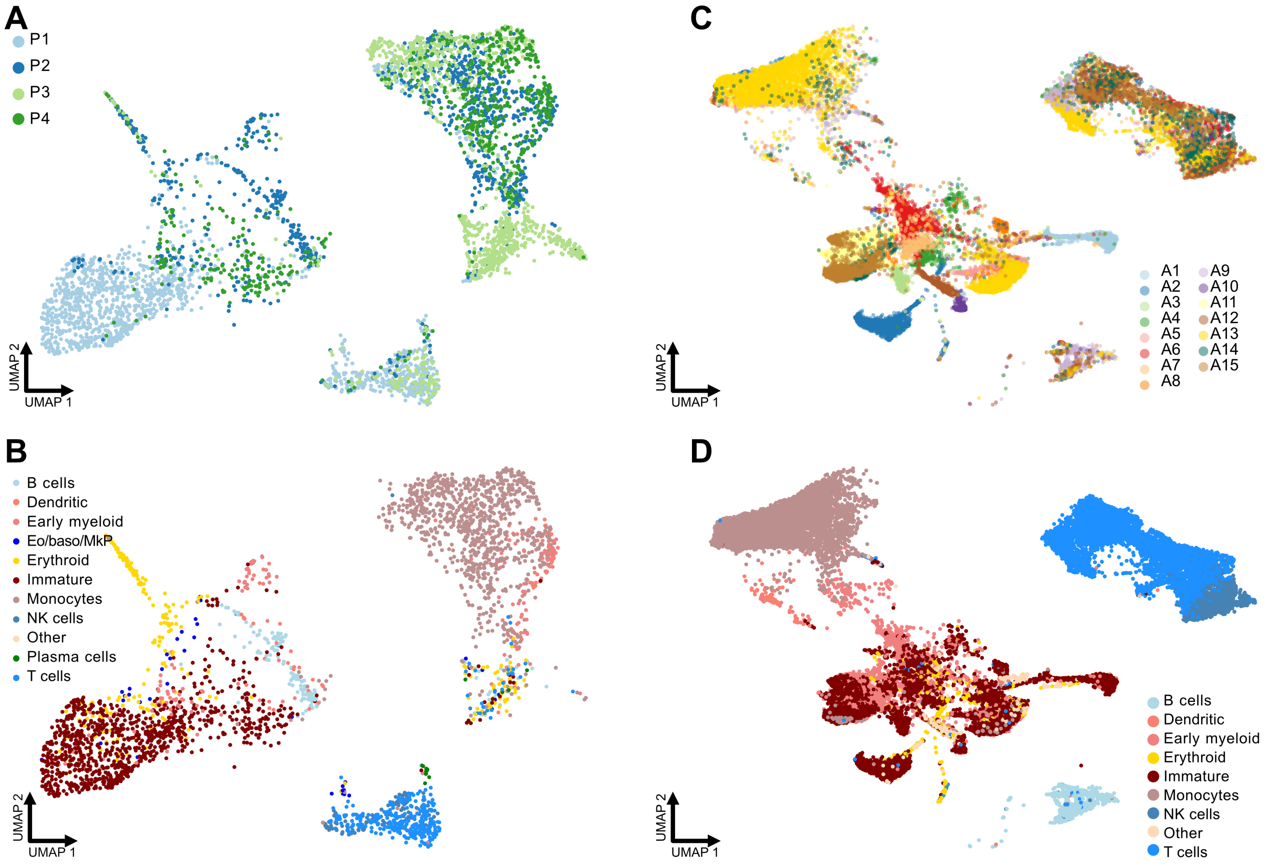

**Figure S2:** UMAP plots showing the patient labels and detailed cell type labels and for the AML Smart-Seq2 dataset [2] respectively in (A) and (B) and the AML 10X patients [3] in (C) and (D). The UMAP were calculated in the original publications. The cell types shown in (D) are extracted from the original publication. For consistency, the cell types labels in (B) were computed in the same way as in (D) [3]. The cells were projected onto a reference atlas of human hematopoiesis [6] as described in [6].

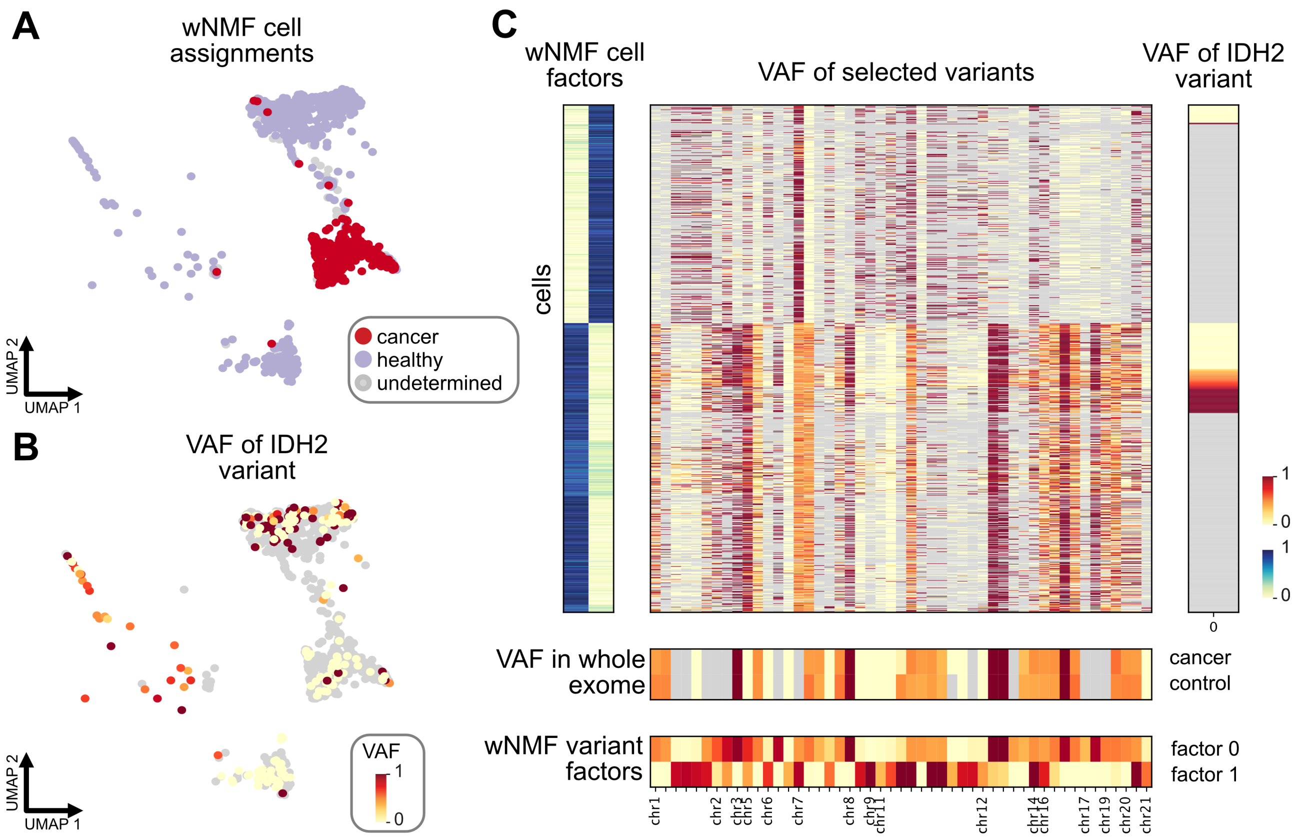

**Figure S3:** (A) UMAP plot showing our cell labels for patient P3 of the AML Smart-Seq2 dataset [2]. (B) UMAP plot showing the VAF of the cancer driving IDH2 somatic variant used to label the cancer cells in the original publication. Cells in grey have insufficient coverage of that variant ($\leq$2 reads). (C) VAF of selected variants for the cells of patient P3. The cells are sorted by cell factors and the subset of variants are selected based on difference between the factors ($\geq$0.3). Grey values have too low coverage ($\leq$ 2 reads for scRNA and $\leq$ 5 for whole exome data). The right-most heatmap shows the VAF of the IDH2 variant, which is found in a subset of the cells of factor 1.

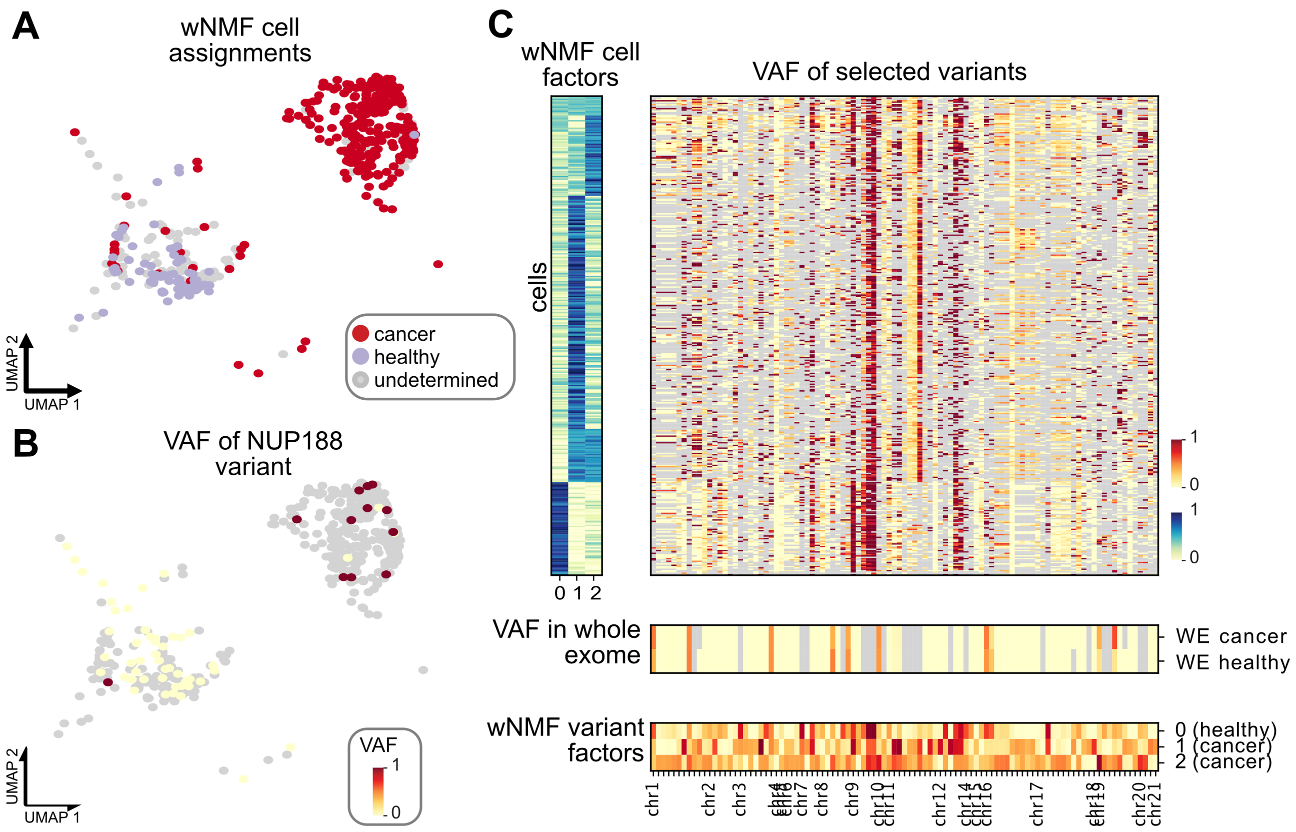

**Figure S4:** (A) UMAP plot showing our cell labels for patient P4 of the AML Smart-Seq2 dataset [2]. (B) UMAP plot showing the VAF of the cancer driving NUP188 somatic variant used to label the cancer cells in the original publication. Cells in grey have no coverage of that variant. (C) VAF of selected variants for the cells of patient P3. The cells are sorted by cell factors and the subset of variants are selected based on difference between the factors ($\geq$0.3). Grey values have too low coverage ($\leq$ 2 reads for scRNA and $\leq$ 5 for whole exome data).

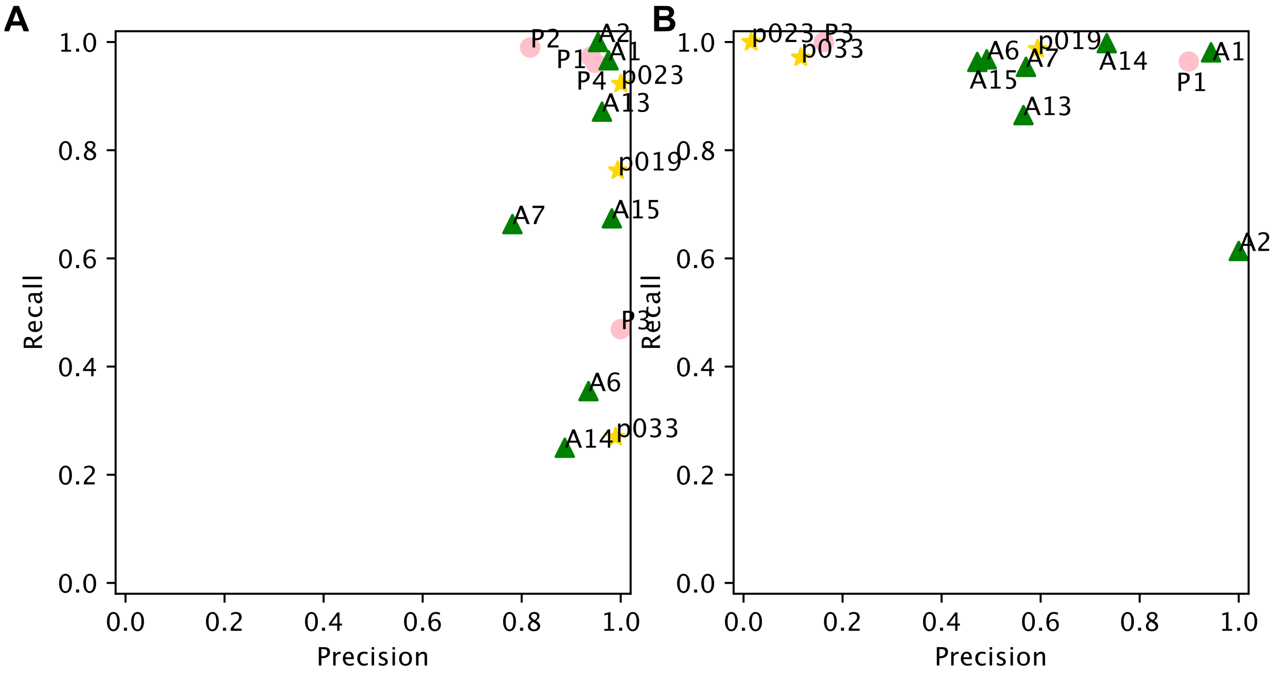

**Figure S5:** We compare our cancer cell labels to a reference cell labels provided by the cell types. In (A) all Blasts for [2], Early myeloid, Erythroid, Immature or Monocytes for [3] and all Tumor cells for [4] are called cancer and compared to our cancer cell assignments. Patients with lower recall are missing some cancer cells and our labels are potentially catching only a subclone of the full cancer population. In (B), all T/NK cells for [2], B cells, NK cells or T cells for [3] and all Ciliated or Club cells for [4] are called healthy and compared to our healthy cell assignments. We expect more cells than only these cell types to be healthy cells, which explains the lower precision for some patients.

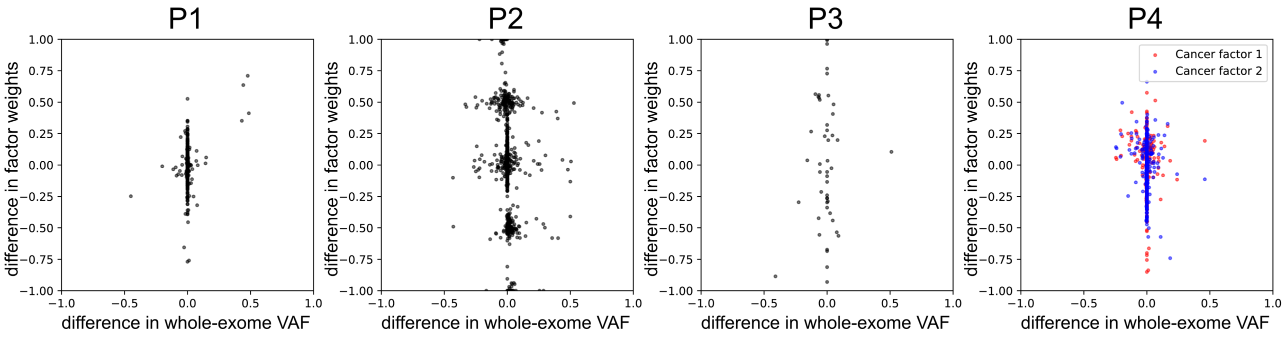

**Figure S6:** We compare the difference in variant allele frequency between the cancer and healthy whole-exome sample on the x-axis ($VAF_{WE:cancer} - VAF_{WE:healthy}$), to the difference in weight between the cancer and healthy factors on the y-axis ($V_{cancer} - V_{healthy}$). One point represents one variant. We show only variants that have sufficient coverage in all wNMF cell factors (>20% of cells covered) and in the whole-exome data (≥5 reads found both in healthy and cancer). Points enriched in the whole-exome cancer versus healthy will have higher values on the x-axis and points enriched in the cancer factors will have higher values on the y-axis. For patient P4 we found two cancer factors, each shown in one color.

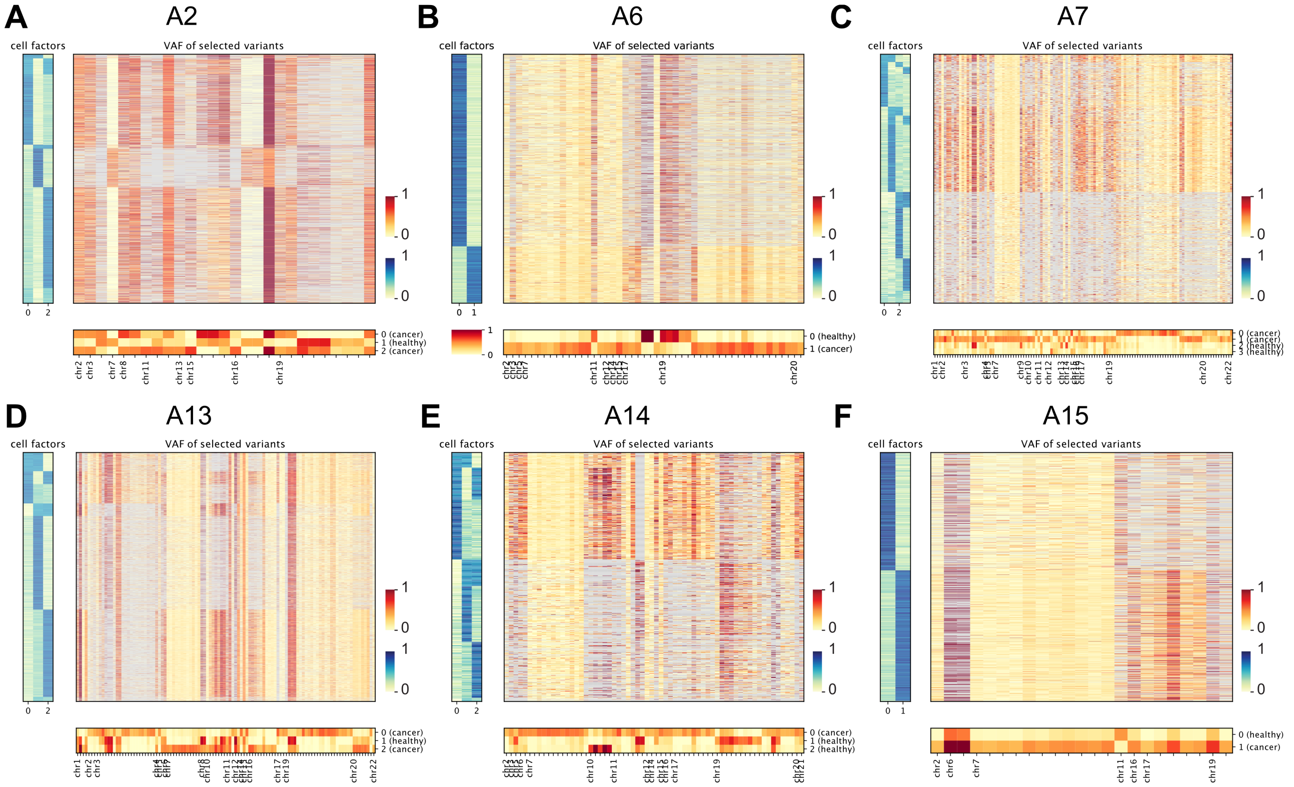

**Figure S7:** VAF of selected variants for all AML 10X patients where CCLONE succeeds in finding clones [3] (excluding A1 which is shown in main Figure 3B). The cells are sorted by cell factors and the subset of variants are selected based on difference between the factors ($\geq$0.3). Grey values have coverage $\leq$ 2 reads.

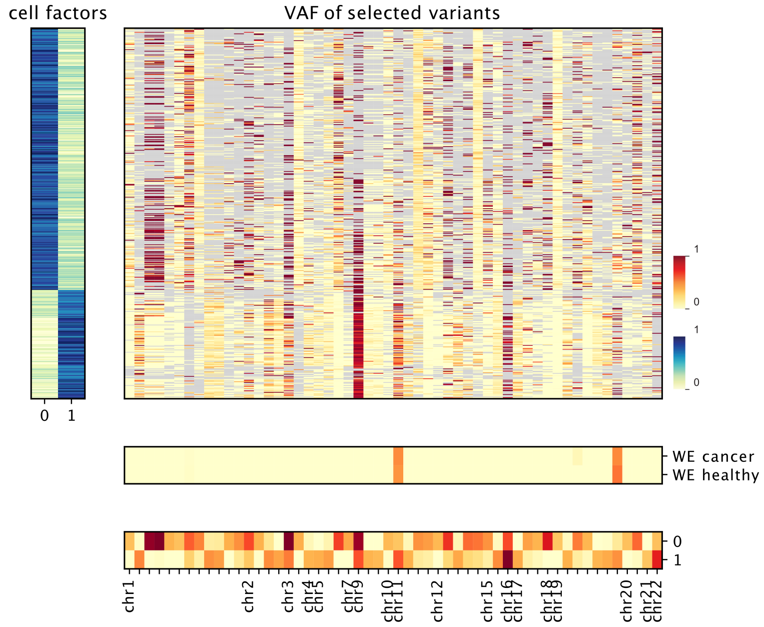

**Figure S8:** VAF of selected variants for P2 excluding donor cells. The cells are sorted by cell factors and the subset of variants are selected based on difference between the factors ($\geq$0.3). Grey values have coverage $\leq$ 2 reads.

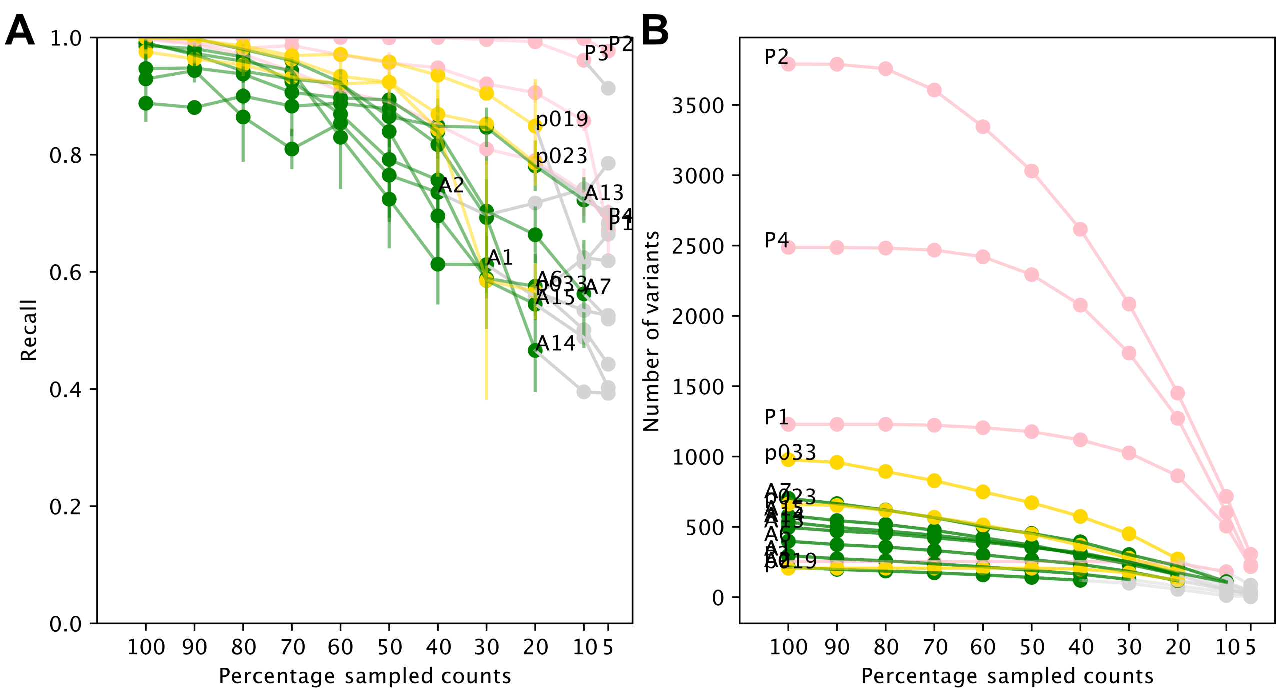

**Figure S9:** For every patient we randomly subsample counts from the reference and alternative count matrices, and rerun the wNMF on the subsets. Comparing the cancer cell labels between the full dataset and the subset, we get the recall as a function of the percentage of sampled counts in (A). Figure (B) shows the number of variants still present in the data as a function of the percentage of sampled counts. The values are plotted in grey if the number of variants with sufficient coverage is lower than 100.

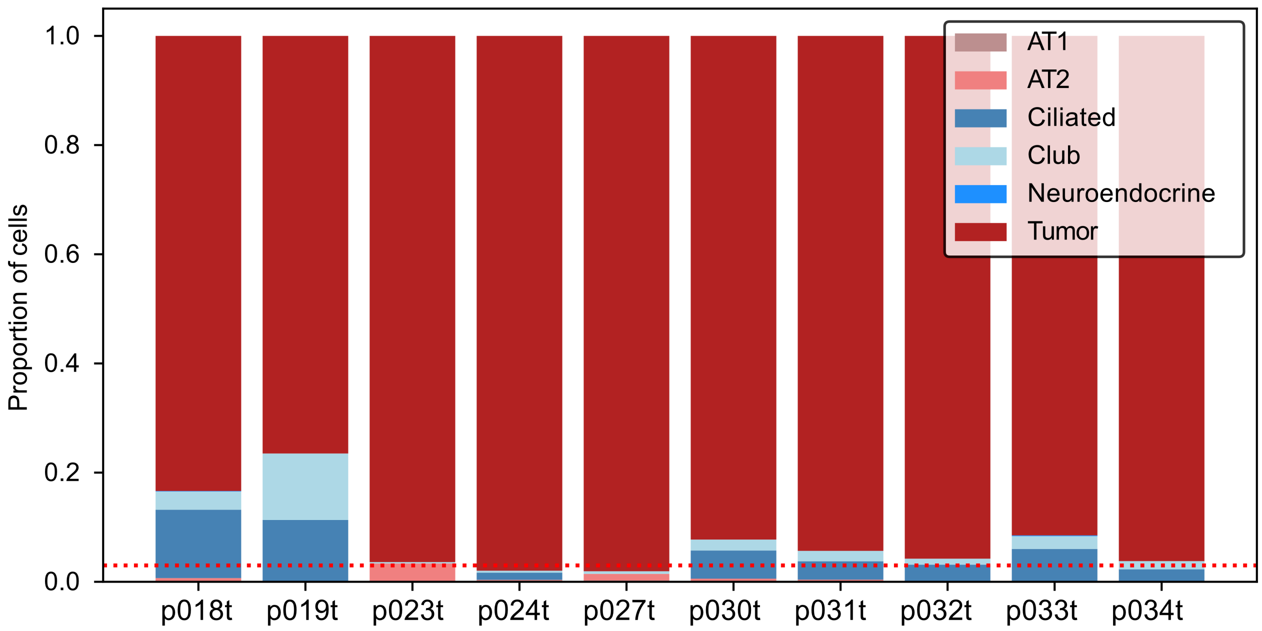

**Figure S10:** Proportion of cells assigned to each cell type for the tumor sample of each patient. The dotted line shows the threshold used for patient filtering. p024 and p027 were excluded because they have too few non-tumor cells in the tumor sample. p034 was filtered out due to the very low total number of cells (132 cells).

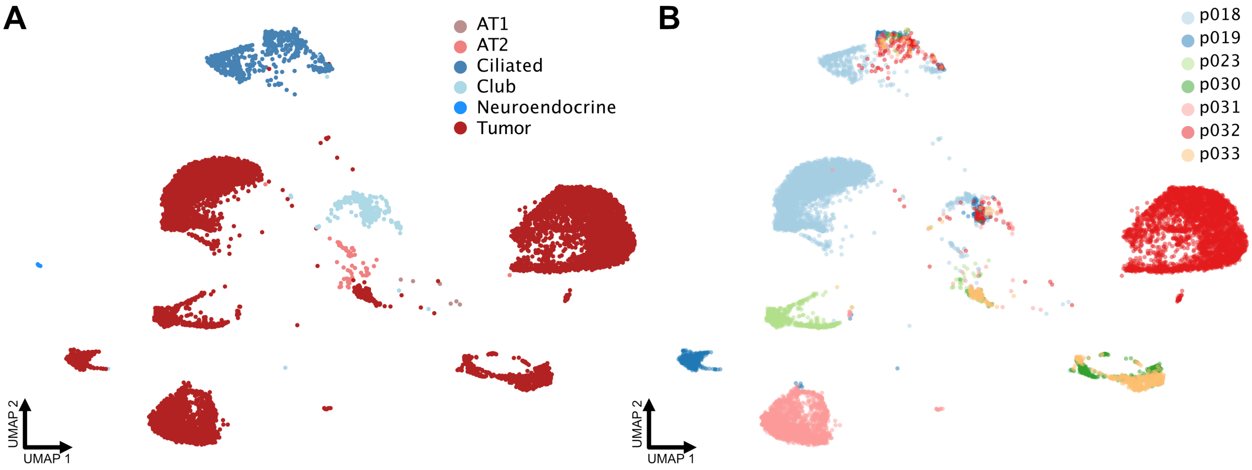

**Figure S11:** UMAP plots showing the detailed cell type label and patient labels for the lung adenocarcinoma dataset [4] respectively in (A) and (B). The UMAP and cell type labels were calculated in the original publications.

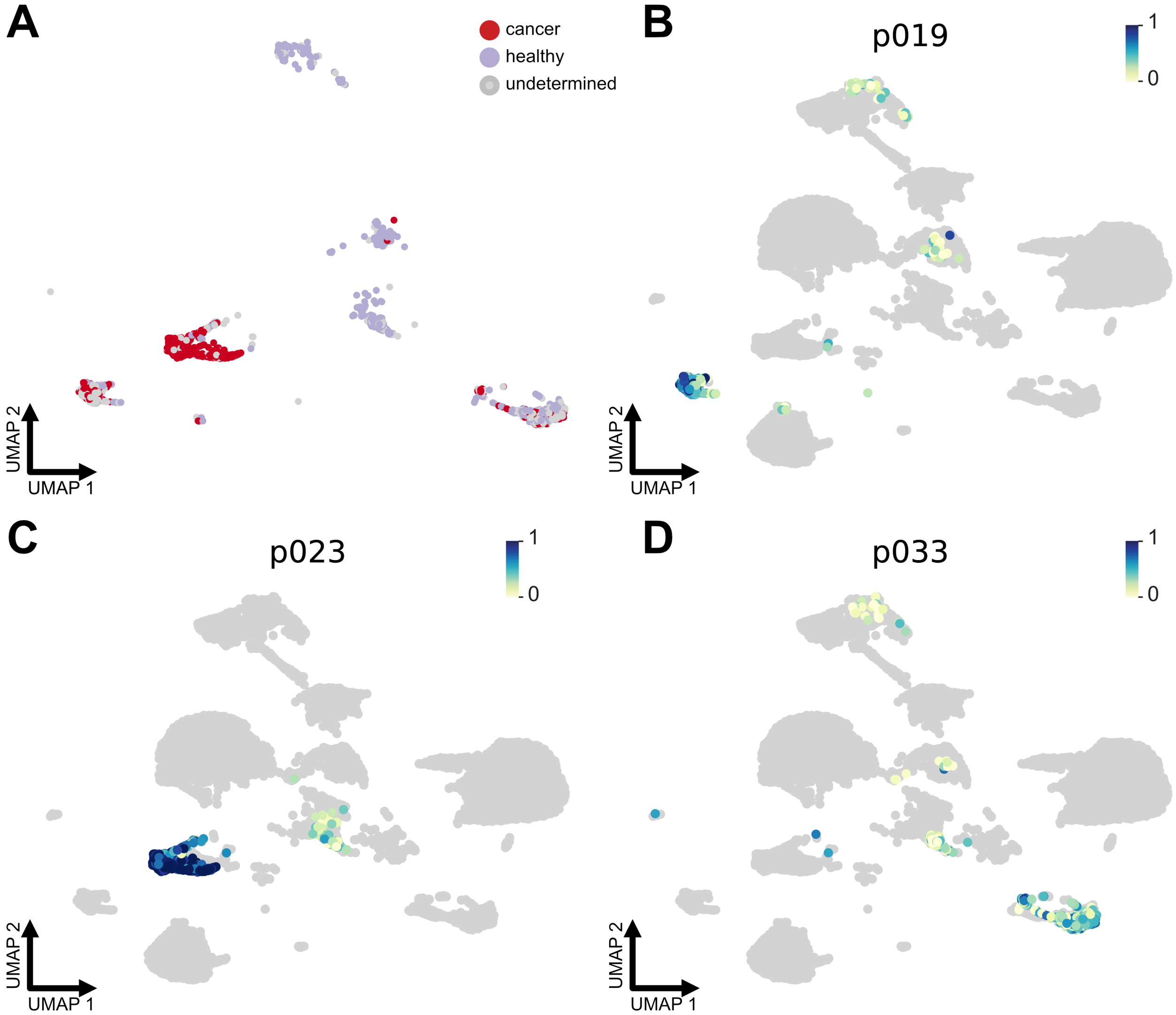

**Figure S12:** UMAP plots showing the CCLONE cancer cell label for the lung adenocarcinoma dataset [4] for patients p019, p023 and p033 in (A) and the weights of the cancer factors for patient p019 in (B), p023 in (C) and p033 in (D)

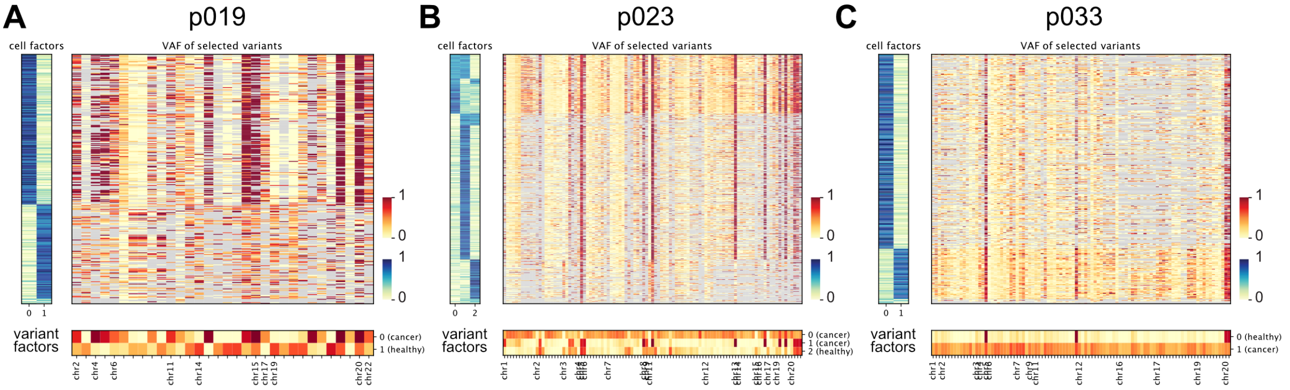

**Figure S13:** VAF of selected variants for p019 in (A), p023 (B) and p033 in (C) for the lung adenocarcinoma dataset [4]. The cells are sorted by cell factors and the subset of variants are selected based on difference between the factors (≥0.3). Grey values have coverage ≤ 2 reads.

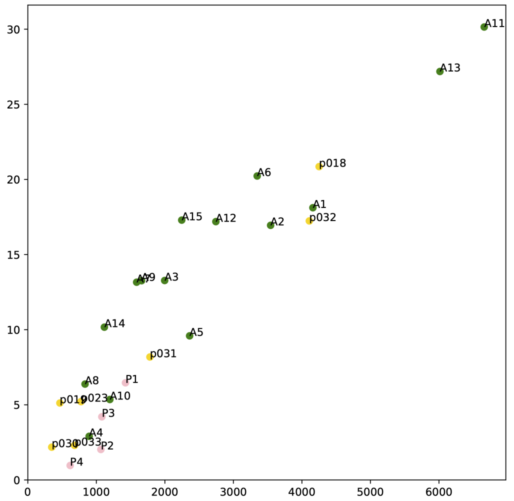

**Figure S14:** Overview of the runtime for variant calling for chromosome 1 with Cellsnp-lite on a Dual Xeon E5-2650v2 (8cores/2.6GHz) and 15 GB of memory for all patients analysed in this work. The variant calling was performed for all chromosomes in parallel and chromosome 1

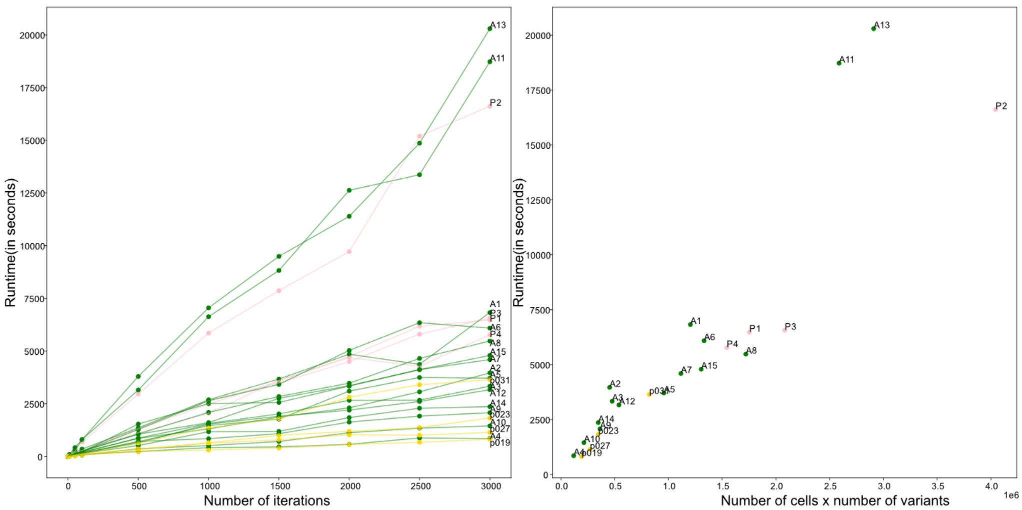

**Figure S15:** Runtime of the wNMF as a function of the number of EM iteration in (A), and of the size of the input variant call matrices (number of cells times number of variants) in (B).

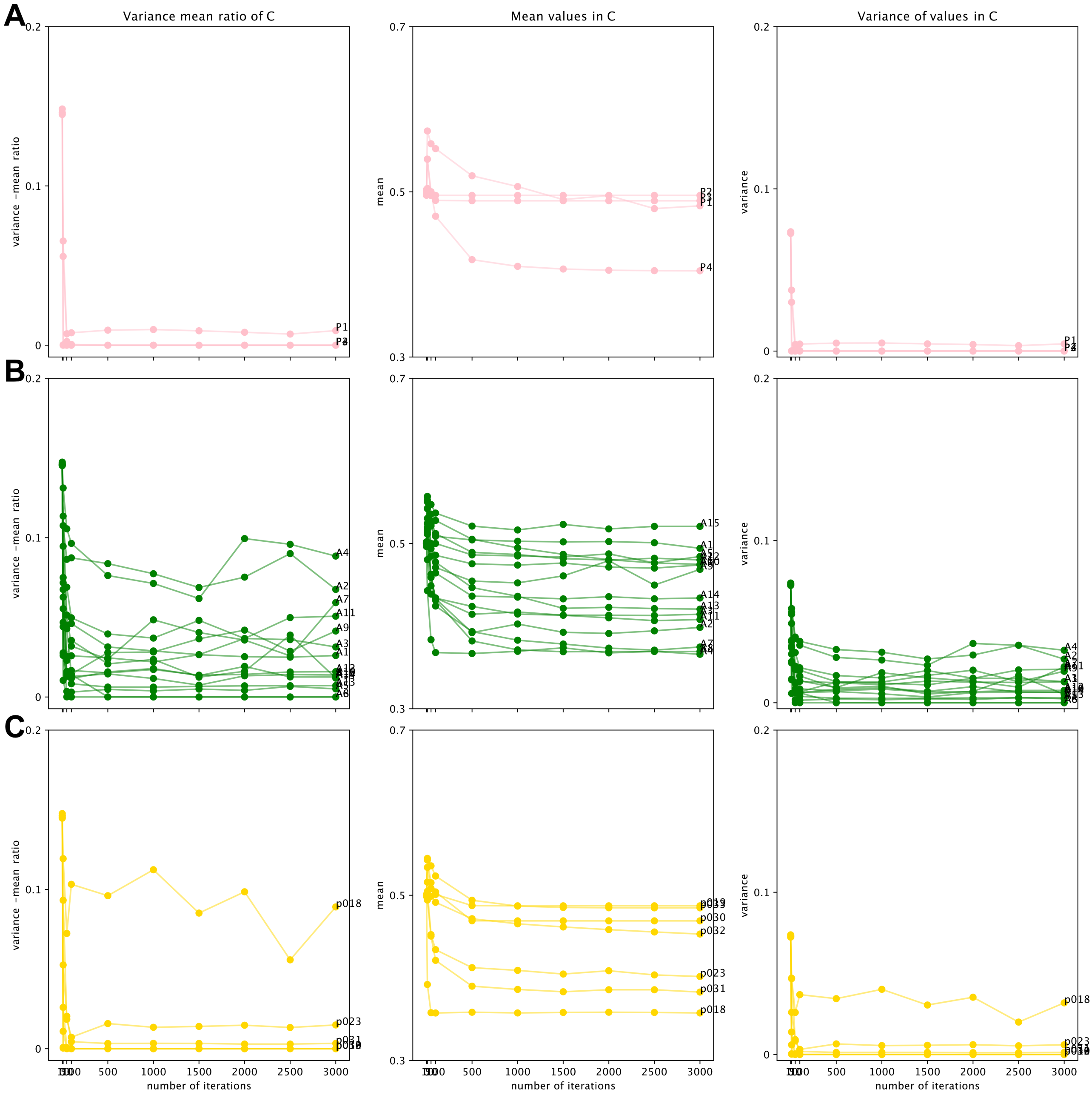

**Figure S16:** We compare the mean and variance of all values in the C matrix over multiple randomly initialised runs of the wNMF as a function of the number of EM iterations. We show this result for the AML Smart-Seq2 dataset [2] in (A), the AML 10X patients [3] in (B) and the lung adenocarcinoma [4] patient in (C). The variance between multiple runs quickly decreases after less than 500 EM iterations. Overall the variance between multiple runs is significantly lower than the mean values in the matrices, indicating that the wNMF is robust with respect to the initialisation.
